# Level of completion along continuum of care for maternal and child health services and factors associated with it among women in Arba Minch Zuria Woreda, Gamo Zone, Southern Ethiopia: a community based cross-sectional study

**DOI:** 10.1101/735456

**Authors:** Dereje Haile, Mekdes Kondale, Eshetu Andarge, Abayneh Tunje, Teshale Fikadu, Nigussie Boti

## Abstract

**Background:** Completion along continuum of care for maternal and newborn health(MNH) service like antenatal care, skilled birth attendance and postnatal care services is one of the currently recommended strategies to reduce both maternal and neonatal mortality to achieve the global target of ending preventable maternal and under five children’s mortality. Although studies on factors affecting each segment of MNH services were well documented in Ethiopia, there is a dearth of evidence on the level of continuum of care and factors associated with it. Thus, this study tries to fill this gap in the country in general and in the study area in particular.

**Methods:** A community-based cross sectional study was conducted among 438 postnatal women who gave births in the last one year in Arba Minch Health and Demographic Surveillance Site. The sample women were selected by using computer generated random numbers from the list of women who gone at least six-weeks after birth. A pre-tested structured interviewer-administered questionnaire was used for data collection. Data was entered and coded in Epi-data and analysed using SPSS software version 23. Binary logistic regression model was fitted to identify factors associated with the outcome variable.

**Results:** The overall completion along the continuum of care was 42(9.7%). The factors significantly associated with continuum of care completion were early antenatal booking (before 16 weeks) [AOR: 10.751, CI (5.095, 22.688], birth preparedness and complication readiness [AOR: 2.934, CI (1.414, 6.087), pre-pregnancy contraception utilization [AOR: 3.963, CI: 1.429,

10.990], employed women [AOR: 2.586, CI: ((1.245, 5.371))], and planned pregnancy [AOR: 3.494 CI :(1.068, 11.425)].

**Conclusion:** Completion along continuum of care was low in the study area. Thus, efforts in improving completion of the cares should focus on early booking during antenatal period, reducing unplanned pregnancy, and improvement on birth preparedness and complication readiness interventions.

## INTRODUCTION

Maternal morbidity and mortality is a major public health problem with high discrepancy between high and low income countries[1, 2]. Worldwide, according to WHO 2015 report, 99% of maternal deaths occurred in developing countries and out of this, around 66% deaths occurred in Sub-Saharan Africa[1]. The MMR and life time risk of maternal mortality was 20 and 27 times higher in developing countries when compared to developed ones. Similarly in Ethiopia, despite the progressive increase in coverage of MNCH service in the last 25 years, the 2016 EDHS report revealed that high rate of maternal mortality ratio (412 maternal deaths per 1000000 live births)[3]. However, nearly 75% of all maternal deaths are due to direct obstetric causes and preventable with highly cost effective interventions[4]. Sub-Saharan Africa and South east Asia regions account for 80% of total neonatal deaths worldwide. A child born in these regions is nine times more likely to die in the first month than a child in a high-income country [5]. According to the 2017 UNICEF report, Ethiopia ranked as one of the five countries which accounted for half of all newborn deaths in the world with India, Pakistan, Nigeria, the Democratic Republic of the Congo the remaining in the list [6].

Continuum of care (CoC), which usually refers to continuity of individual care and which is one of the health services strategic framework and indicator that measure coverage of maternal, newborn, child and adolescents’ health (MNCAH) is currently given attention[7]. It has two dimensions: time and place. CoC at its time dimension refers to a situation where a woman and her child receive maternal, newborn and child health (MNCH) services from pre-pregnancy, pregnancy, childbirth, and postpartum to childhood. The place dimension focuses on integration among household level, community-level and facility-level MNCH care and referral to advanced-level care if needed[8, 9].

Completion along continuum of care for maternal and newborn health service like antenatal care, delivery attended by skilled personnel and care after child birth services are one of the strategy to reduce both maternal and neonatal mortality[9]. Furthermore, improvement in completion along continuum of care for MNCH service or life course approach was one of the recommendations to achieve the global target of reduction of maternal mortality to 70 maternal death per 100,000 live birth and ending preventable under five children’s mortality[10]. The completion in the continuum of care has outweighing advantages over separate cares or intervention because each stage of continuum of care builds on the success of the previous stages [11-16]. However, evidences from the globe suggest that women completing the continuum of care for maternal and child health services are very low and women do not access MNCH services serially along continuum of care [17-19].

Ethiopia with a remarkably low progress in the successive EDHS reports in each segment of the MNH cares [3] may not be different in utilization of MNCH services along the continuum of care. However, to the best of our knowledge, there is a dearth of documented evidence on the magnitude of completion of the services along continuum and factors associated with completion in the country in general and in our study area in particular. Thus, to respond to these needs of study and to fill this gap in the scientific literature, our study aims to assess the level of completion along continuum of care for MNH services and identify factors associated with it among the target women in Arba Minch Health and Demographic Surveillance Site, Southern Ethiopia. Given the high toll of maternal and newborn morbidity and mortality in Ethiopia, the findings of this study will be of paramount importance for policymaking and programme planning.

## Methods

### Study setting and design

The study was conducted in Arba Minch Health and Demographic Surveillance Site, which is located in Arba Minch Zuria Woreda, Gamo Zone, Southern Ethiopia, 454 kms to the South of Addis Ababa, the capital city of Ethiopia. The surveillance site consists of eight rural and one semi-urban kebeles) which were selected in representation of all kebeles in the district. [*Woreda: an administrative unit corresponding to district in other parts of the world* and *Kebele: the smallest administrative unit in the current Ethiopian government structure under woreda].*

In Arba Minch HDSS, there was continues registration and recording of maternal and newborn health services. A community based cross sectional study was conducted from Febraury15/2019 to March 15/2019. All women (n=595) who had registered from December 2017 to December 2018 in Arba Minch HDSS who gave birth after booked for ANC in health facilities and who gone at least six weeks in the postpartum period were considered as study population.

### Populations of the study

The source populations for this study were all women in Arba Minch HDSS who gave birth after booked for ANC in health facilities and were at or after six weeks of childbirth. Those who gave birth after booked for ANC in health facilities and were at or after six weeks of childbirth in the selected kebeles for data collection period were the study populations of the study. Women who reside in the study area for less than six months and who were critically ill at the time of data collection were excluded from the study.

### Sample size Determination

The sample size for this study was calculated using statcalc menu of Epi-info software version 7 initially using the assumptions for single population proportion with estimated prevalence of 50%(because of the absence of study in a similar setting), confidence level of 95% and 5% degree of precision which gives 384. With a consideration of 10% non-response rate, the the final sample size was 423. Secondly, a two-population proportion with consideration of factors affecting continuum of care for MNH care was made. Among the biological plausible factors selected, the largest sample size was obtained using the assumptions of 80% power, 95% confidence level, proportion of completion among women not knowledgeable on pregnancy danger signs(P1=55%) and proportion of completion among women knowledgeable on pregnancy danger signs and 10% non-response rate (P2=69.5%) [20] which gives 438. Since the sample size obtained using two population proportion considerations (n=438) was larger than the sample size for single population proportion (n=423), it was used as the final sample size for the study.

### Sampling technique

The samples were obtained from a secondary data in the HDSS. The maternal health data consists of different health and health related data of women in relation to their pregnancy, labor and delivery and postnatal care. A total of 1316 women who gave live birth in the last one year before the data collection period were registered in the HDSS registry. Of those women, 595 fulfilled the inclusion criteria and a sampling frame of these women was used to select the sample women of this study. The required number of women in the sample for the study (n=438) were selected after proportional allocation of women to each kebele based on the number of these women in the kebeles. Thus, a separate list of women was prepared for each kebele and the required samples were selected using computer generated random numbers (s1fig).

### Data collection methods and procedure

Nine HDSS data collectors who had prior experience in data collection were recruited for data collection and three public health officers from the different district health facilities supervised the data collection. Both data collectors and supervisors were given a daylong intensive training on the data collection methods and instruments. The data was collected from women’s by using semi-structured interviewer administered questionnaires. A pretested, structured, interviewer administered questionnaire was used. The questionnaire was adapted from EDHS and other published literatures. House hold wealth index of the study women was assessed by using a tool adapted from the Ethiopian Demographic and Health Survey 2016 wealth index assessment tool. The supervisors and data collectors accessed houses of the sampled women by the guidance of the local Health Development Army (HDA) leaders in each kebele. The data collectors were given the list of women selected for the interview in their respective kebeles in advance.

### Data quality management

An intensive training was provided to the data collection team by the team of investigators with particular focus on the contents of the questionnaire. A practical role play on interviewing skills was exchanged among the data collection team. The questionnaire used to obtain data from the study women was initially prepared in English and translated to the local language (*Gamotho*) by an expert in the language and finally back translated by another expert to English to check its consistency with the original meaning. Finally, a structured questionnaire in the local language was used for data collection. Before commencing data collection, pre testing was conducted on 5% of the sample size (22 women) out of the study area that had similar characteristics with the study population and was out of the sampled clusters in the study area. In response to the pretest findings, the necessary amendments were made to unclear and confusing questions. To minimize social desirability bias, women were interviewed in a separate private place in their own household compound.

### Data analysis

Data was entered in to Epi-data software version 3.1 and then exported to SPSS version 23 statistical package for analysis. Descriptive statistics was done to quantify level of completion on the continuum of care for MNH services and other explanatory variables. Findings have been summarized in tables and graphs using frequencies, percentages and standard deviations. Wealth status of the individual household was analyzed using principal component analysis method of factor analysis after checking the fulfillment of assumptions using Kaiser-Meyer-Olkin measure of sampling adequacy and Bartlett’s Test of Sphericity. Binary logistic regression model was fitted to identify factors associated with completion along continuum of care. Model fitness was checked by Hosmer and Lemeshow goodness of fitness test and multicollinearity between the explanatory variables was checked using variance inflation factor (VIF>10). Initially, bivariate logistic regression analysis was performed between dependent and each of the independent variables in sequence. Variables having p-value of <0.25 in bivariate logistic regression were selected as candidates for multivariable logistic regression analysis and entered sequentially by using backward stepwise regression method. Association between outcome variable and independent variables reported by using adjusted odds ratio and its 95% CI, and variables having p value less than 0.05 in multivariate logistic regression model were considered as statistically significant.

### Definition and measurement of variables of the study

The outcome variable for this study is completion along continuum of care for maternal and newborn health services (CoC). The completion status has been classified as completed the continuum or have not completed the continuum of care. Women were having completed the continuum when:

1. They have had 4 or more ANC visits by a skilled health personnel (Medical doctors, nurses, midwives, health officers and community health extension workers) and
2. Have had a childbirth aided by Skilled Birth Attendant(SBA) e.g., doctor, nurse, midwife, and health officer at health institution and
3. Have had attended Post Natal Care(PNC) for the mothers and newborns at least once after discharge from health facilities within six weeks by a skilled health personnel(doctor, nurse, midwife and health officer) at the health facility or within the first week by community health extension workers during their home visit.

Women were classified as not completed the continuum of care if they missed any one of the above visits/attendances at any level.[17, 21-23].

### Different groups of explanatory variables have been considered in the study by reviewing related literatures and theories

1. Individual women and household socio-demographic factors considered include woman’s age, education, employment status, partner’s education, household wealth status (*a composite indicator of socio-economic status of women which divides the households into five or four or three categories and were derived using principal component analysis based on information from housing characteristics and ownership of household durable goods*).
2. Women’s related factors considered include exposure to mass media, membership of health insurance, distance to health facilities, means of transportation, women’s autonomy to seek health care decision, knowledge on key pregnancy danger signs.
3. Obstetric factors included birth order, time for first ANC visit, pre pregnancy contraception utilization, desire on pregnancy, selected elements of ANC services provided, and birth preparedness and complication readiness(BPCR).

Autonomy in household decision making: A woman was said to have decision making power on seeking MNH service if she alone or with her husband decide on seeking MNH service[18]. Time for first ANC visit: the time or gestational period at which the pregnant women first attend ANC clinic was classified as in the first trimester or after first trimester[24, 25]. Knowledge on key pregnancy danger signs: women were classified as knowledgeable if they mentioned at least two of the four key danger signs of pregnancy (vaginal bleeding, severe headache, blurring of vision and feet or face swelling) if not they were classified as not knowledgeable[17].

Essential elements of ANC service provided: Women were classified as received full services and not received full services. Full service was measured as when women’s received the seven selected essential antenatal care services (blood pressure measurement, blood sample collection, urine sample collection, tetanus toxoid(TT2+) vaccination, iron foliate(90+) supplementation and health education on danger signs and nutrition). Full services was considered as not received when one of the seven selected elements of antenatal care services was not provided during ANC visit [21, 26]. Means of transportation to health facilities was classified as on foot, by bicycles, or by car[18, 19]. Women were considered as well prepared for birth and its complications when they reported that they have implemented five or more components of BPCR otherwise she was considered as ‘not well prepared’. The components of BPCR considered in this study were identified place for birth, identified birth attendants, saved money, identified emergency transportation, identified labor and birth companion, identified blood donors as needed, and identified care giver to children’s at home when a mother was away [27, 28]. Pre-pregnancy contraception utilization was measured by women’s report whether they used any modern contraception method before their recent pregnancy and birth. Desire on pregnancy measured as planned or unplanned. Planning for the last pregnancy was measured by asking women whether the last pregnancy occurred when she had desired additional children or not. Thus, unplanned pregnancy was considered when women had no desire or no more desire for additional children or pregnancy occurred earlier than the desired time. Exposure to mass media was measured by asking (Yes/No question) on women’s habit of watching TV or listening to radio to access relevant information on maternal and child health services.

### Ethical approval and consent to participate

Ethical clearance was obtained from the Institutional Research Ethics Review Board of Arba Minch University. Respondents were informed about the purpose and procedure of the study and oral consent was obtained from participants. The privacy and confidentiality of the information given by each respondent was kept confidential and anonymous. Formal letter of permission was obtained from the local health offices.

## Result

In this study, four hundred thirty two women were volunteered to give information making a response rate of 98%.

### Socio-demographic characteristics of the study participants

The median age of the respondents were 27.5(IQR=5), and 416(96.2 %) were married. More than two-third (69%) of them were followers of the protestant religion One hundred seventy three (40.04%) of the study women were unable to read and write while 180(43.68%) of their husbands has got a formal education to primary level of education ((Table 1).

**Table 1:**
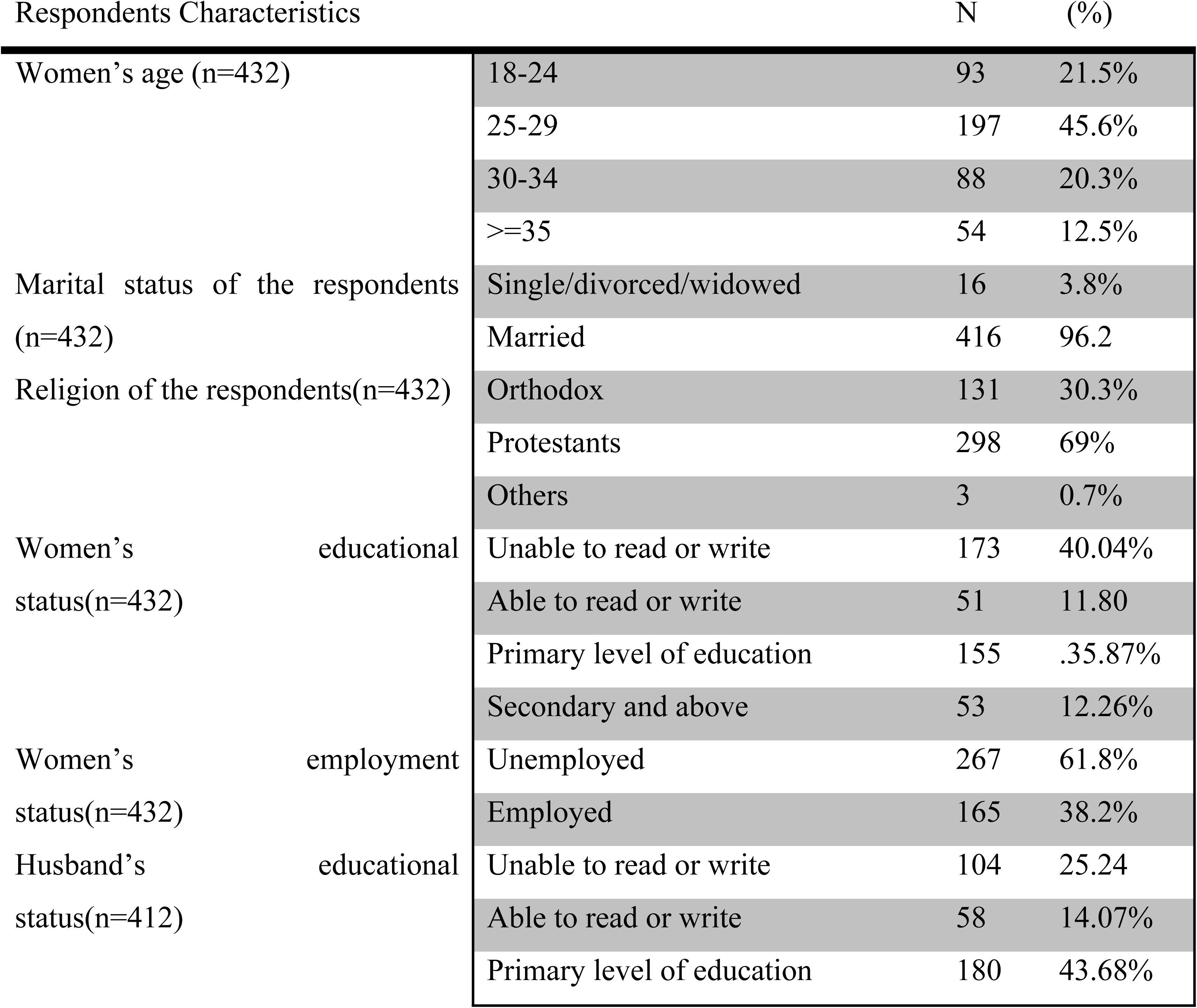
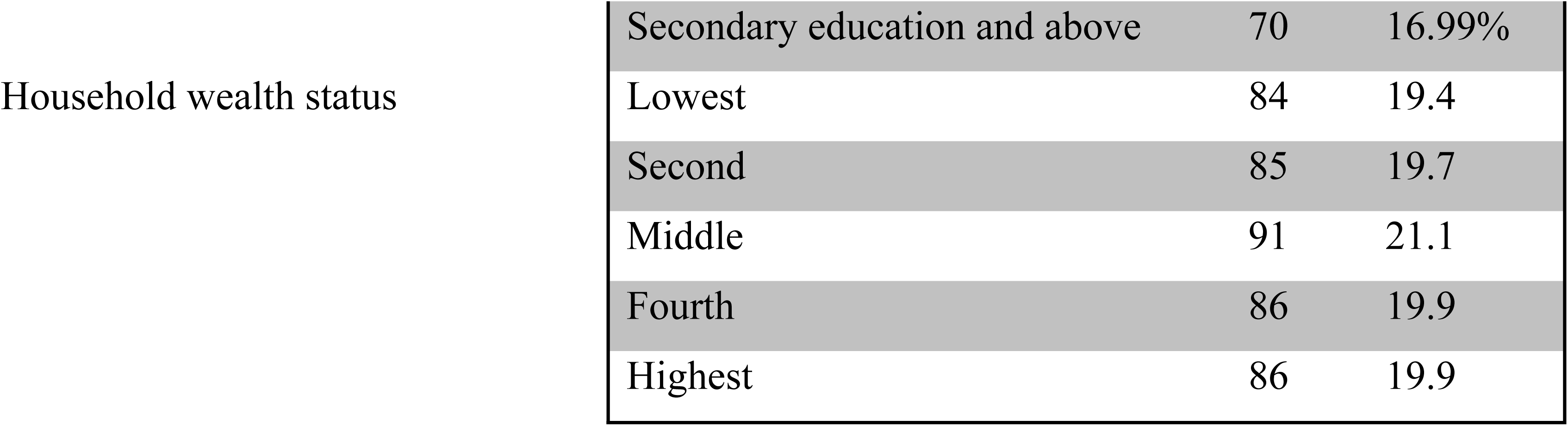
Socio-demographic characteristics of postpartum women in Arba Minch Health and Demographic Surveillance Site, Feb 15-March 15/2019.

### Women health service access and behavior related characteristics

Regarding accessibility of maternal and newborn health care services, more than half (58.3%) reported that the average time to reach health facilities was less than thirty minutes and the majority (83.1%) of participants travel on foot to health care services. More than half 222(51.4%) of respondents were insured by community based health insurance. Majority of the study women (79.6%) make decisions for maternity care autonomously (Table 2).

**Table 2:** Women related characteristics among postpartum women in Arba Minch HDSS, Feb 15-March 15, 2019.

| Respondents Characteristics |  | Numbers(n) | (%) |
| --- | --- | --- | --- |
| Means of transportation to health facilities(n=432) | By motorcycle/car | 73 | 16.9% |
|  | On foot | 359 | 83.1% |
| Perceived required time to reach health facilities(n=432) | < 30 minutes | 252 | 58.3% |
|  | >=30 minutes | 180 | 41.7% |
| Exposure to mass media (radio/TV)(n=432) | Yes | 240 | 55.6% |
|  | No | 192 | 44.4% |
| Frequency of exposure to mass media(n=240) | Always | 65 | 27.1% |
|  | More than once a week | 60 | 25% |
|  | Once a week | 115 | 47.9% |
| Membership of community based health insurance(CBHI) (n=432) | Yes | 222 | 51.4% |
|  | No | 210 | 48.6% |
| Women's autonomy to maternity care (n=432) | Autonomous | 344 | 79.6% |
|  | Non-autonomous | 88 | 20.4% |

### Obstetric characteristics of the study participants

Regarding to obstetric history of respondents, 359(73%) were multiparous, 99(22.9%) were unplanned for pregnancy. More than two-third of the study women were not knowledgeable to pregnancy danger signs (Table 3).

**Table 3:** Obstetrics characteristics of postpartum women in Arba Minch HDSS, Feb 15-March 15, 2019

| Respondent's characteristics | Category | Frequency | % |
| --- | --- | --- | --- |
| Pre-pregnancy utilization of contraception(any modern methods) (n=432) | Yes | 294 | 68 |
|  | No | 138 | 32 |
| Women's knowledge on key pregnancy danger signs (n=432) | Not-knowledgeable | 261 | 60.42 |
|  | Knowledgeable | 171 | 39.58 |
| Birth order(n=432) | First | 73 | 16.9 |
|  | Second | 95 | 22 |
|  | Third | 121 | 28 |
|  | Four and above | 143 | 33.1 |
| Women's desire on recent pregnancy(n=432): | Planned | 333 | 77.1 |
|  |  | 99 | 22.9 |
|  | Not Planned |  |  |
| Time for first ANC booking (n=432) | At or after 16 wks. | 337 | 78 |
|  | Before 16 wks. | 95 | 22 |
| BPCR(n=432) | Not well prepared | 301 | 69.67 |
|  |  | 131 | 30.33 |
|  | Well prepared |  |  |

### Continuum of care for maternal and newborn health services

#### Level of completion along continuum of care

Overall prevalence of complete continuum of care among women who gave birth the last one year after booked for ANC in the health facilities were 42(9.7%)[95% CI: 6.9, 12.5] (Figure 1).

**Figure 1:**
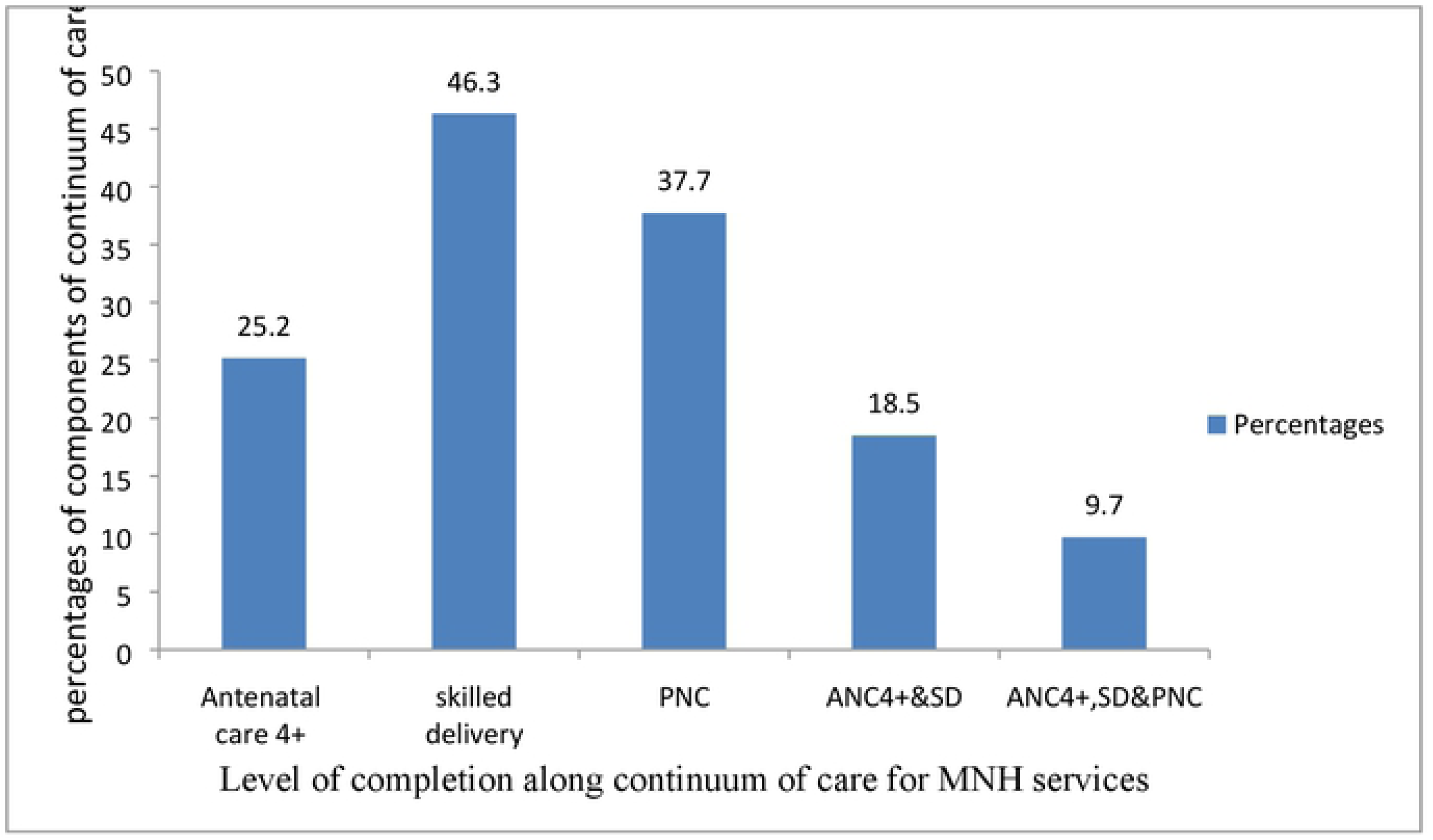
Completion along continuum of care for maternal and child health services among women in Arba Minch Demographic and Health Surveillance site, Feb 15-March 15, 2019.

##### Completion along continuum of care during ante-partum period/pregnancy

Out of all respondents, only 109(25.2%) women completed continuum of care during ante-partum period. Regarding the services provided during ante-partum period, among all respondents 95(22%) completed seven selected essential antenatal services (blood pressure measurement, blood sample examination, and urine sample examination at least once, vaccination for tetanus, HIV testing, provision of information on danger signs and nutrition, and iron provision)(Figure 2)

**Fig 2:**
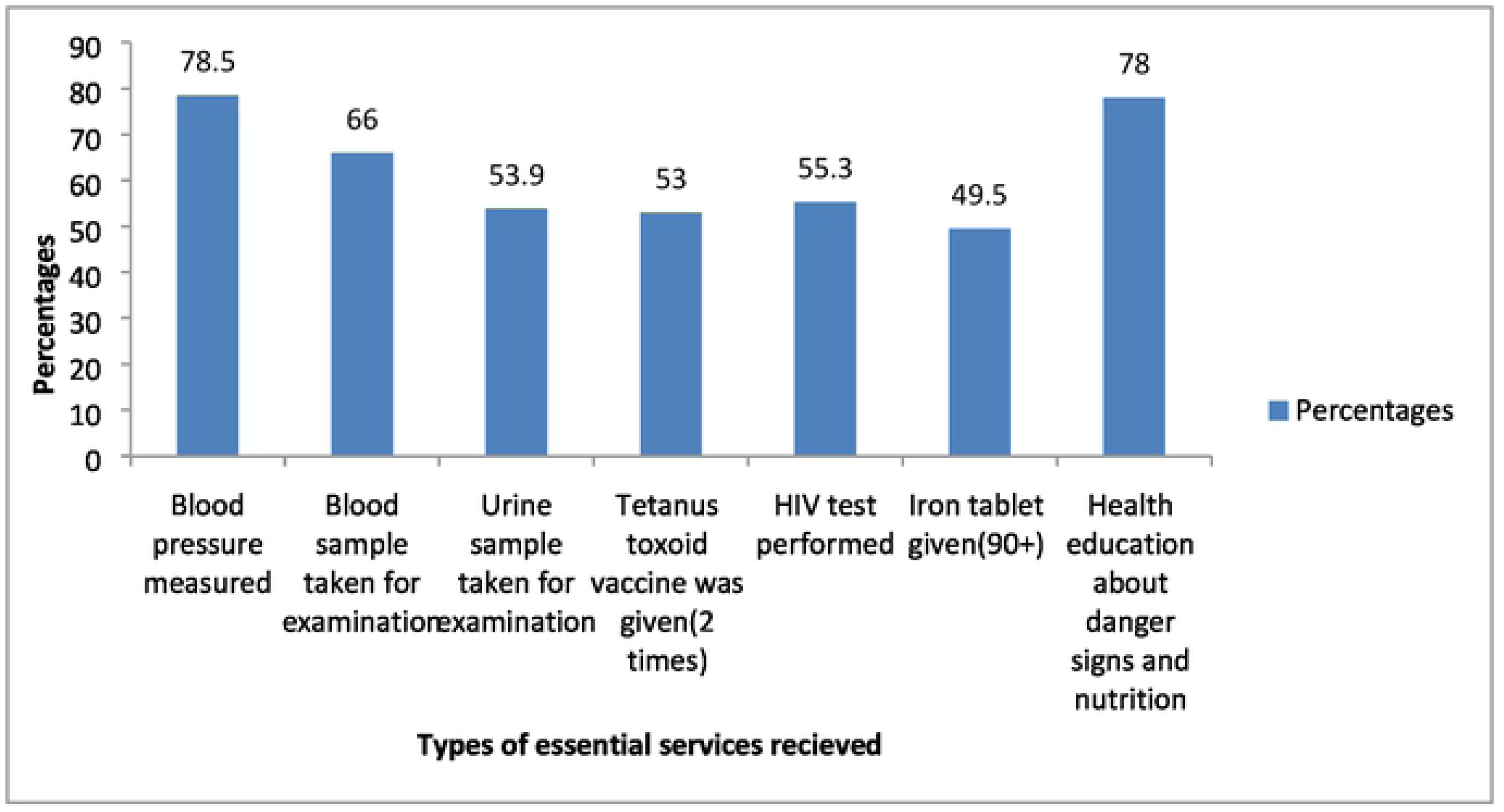
Percentages of postpartum women who received selected essential ANC services during the antenatal period in Arba Minch HDSS, Feb 15-March 15, 2019. Regarding to content of care, among women’s who completed continuum of care, 88.1% received blood pressure measurement. In contrast, 77.4% of respondents who were not completed CoC received blood pressure measurement. The other elements have also showed a similar trend (Figure 3).

**Fig 3:**
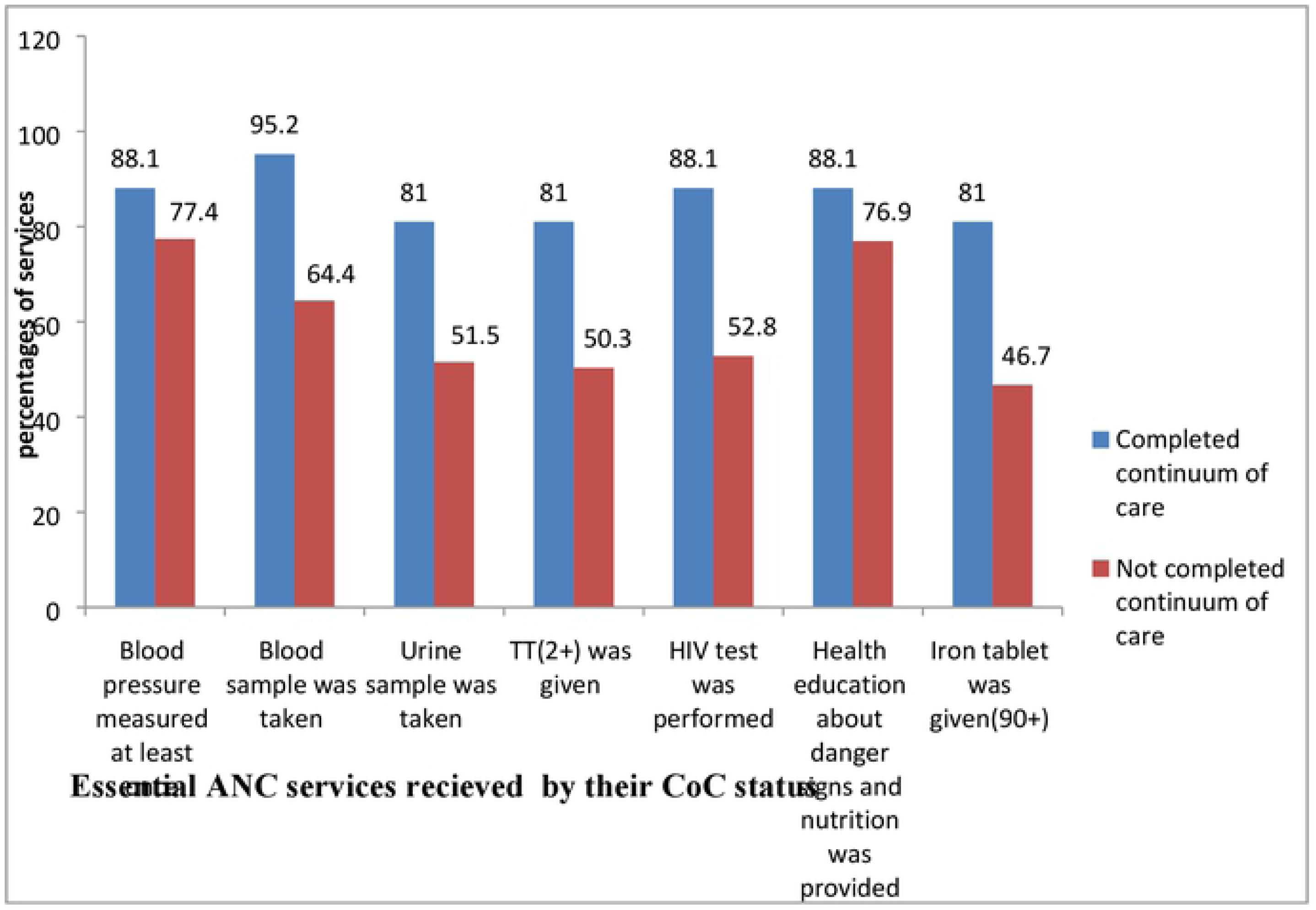
Percentages of selected essential ANC services by their CoC completion status among post natal women in Arba Minch Demographic and Health Surveillance site, Feb 15-March 15, 2019.

##### Completion along continuum of care during intra-partum period

Out of 109 women who completed continuum of care during ante-partum period, 80(73.4%) continued care during intra-partum period. This accounts only 18.5% of the total number of respondents. Regarding maternal and newborn health services provided during inter-partum period, among individuals who had completion of continuum of care during inter-partum period 75(93.8%) had exercised skin to skin contact, and only 4(5%) utilized immediate postpartum contraception during intra-partum period (Table 4)

**Table 4:** Content of care provided to women during intra-partum period in Arba Minch HDSS, Feb 15-March 15, 2019.

| **Newborn services during inter-partum period (n=80)** | **Frequency** | **Percentages** |
| --- | --- | --- |
| **Newborns had got weight measurement** | 73 | 91.3 |
| **Exercised skin to skin contact** | 75 | 93.8 |
| **Initiated breast feeding within one hours** | 53 | 66.3 |
| **Had got necessary immunization** | 62 | 77.5 |
| **Cord care** | 190 | 44.5 |
| **Maternal services during inter-partum period (n=80)** |  |  |
| **Blood pressure measured** | 55 | 68.8 |
| **Counseled about postpartum contraception utilization** | 63 | 78.8 |
| **Immediate postpartum contraception utilization** | 4 | 5 |
| **Counseled about postpartum complication** | 67 | 83.8 |

##### Completion along continuum of care during postpartum period

Regarding to postpartum period, out of 80 women’s who have completed continuum of care during intra-partum period, 42(52.5%) of them continued care during postpartum period and this figure is only 9.7% out of the total number of respondents.

Among respondents who have had completed the continuum of care at postpartum period, after discharge from health facilities, 14(33.3%) had their first PNC contact within two days, 19(45.2%) attended PNC in 3-7 days, 6(14.3%) attended PNC in seven days to two weeks, and 3(7.1%) attended PNC in two weeks to six weeks postpartum. Regarding women’s place of postnatal care, 5(11.9%) attended at hospital, 8(19%) attended at health center, 11(26.2%) attended at health post, and 18(42.9%) received the care at their own home. As regards the contents of care, 28(66.7%) of those completed the CoC have had their children received cord care, and 10(23.8%) have been provided with iron tablets (Table 5).

**Table 5:** Contents of care provided to women during postpartum period in Arba Minch HDSS, Feb 15-March 15, 2019.

| **Newborn care services provided** | **Frequency** | **Percentages** |
| --- | --- | --- |
| **Cord care with chlorhexidine** | 28 | 66.7 |
| **Weight measurement** | 20 | 47.6 |
| **Counseling on infant feeding** | 25 | 59.5 |
| **Immunization** | 27 | 64.3 |
| **Maternal health care services provided** |  |  |
| **Women's counseled about postpartum contraception** | 39 | 92.9 |
| **Women's have utilized postpartum contraception,** | 18 | 42.9 |
| **Women's who were counseled about postpartum complication** | 22 | 52.4 |
| **Women's who were provided with iron tablet** | 10 | 23.8 |

### Factors associated with completion along continuum of care for maternal and newborn health services among postnatal women in Arba Minch HDSS

Multivariable analysis identified five independent factors affecting the completion status of MNCH services. These were time for antenatal booking, pre-pregnancy utilization of contraception, maternal employment, birth preparedness and complication readiness, and pregnancy desire.

The odds of completing continuum of care for maternal and newborn health services was about 11 times higher for those who booked for antenatal care early(before 16 weeks) than their counterparts[AOR: 10.75, CI (5.09,22.68]. Similarly, the odd of completion along continuum of care was higher among women well prepared for birth and its complication during pregnancy than those who were not well prepared [AOR: 2.93, CI (1.41, 6.08). Use of complete continuum of care for MNH services was significantly associated with, respondents who used contraception before their recent birth. The odds of completion along continuum of care for maternal and newborn health services were 4 times higher for those who used pre-pregnancy contraception when compared to their counterparts [AOR: 3.96, CI: 1.4,10.99] and employed women were had also higher odds of completion than non-employed women [AOR: 2.58, CI: ((1.24,5.37))]. Women who planned for their recent pregnancy had also a higher odds of completion as compared to those who did not plan for their recent pregnancy [AOR:3.49 CI:(1.10,11.42)] (Table 6).

**Table 6:** Factors associated with completion of the continuum of care for maternal and newborn health services among postnatal women in Arba Minch Demographic and Health Surveillance Site, Feb 15-March 15, 2019.

| Variables | CoC for MNH services |  |  |  |
| --- | --- | --- | --- | --- |
|  | Yes (%) | No (%) | COR(95% CI) | AOR(95% CI) |
| **Women's education status(n=432)** |  |  |  |  |
| Unable to read or write | 11(6.4) | 162(93.6) | 1 | 1 |
| Able to read or write | 6(11.8) | 45(88.2) | 1.96(0.68,5.60) | 1.931(0.54,6.95) |
| Primary level of education | 17(11) | 138(89) | 1.81(0.82,4.00) | 1.78(0.59,5.36) |
| Secondary and above | 8(15.1) | 45(84.9) | 2.62(0.99,6.89) | 1.93(0.48,7.65) |
| **Women's employment status(n=432)** |  |  |  |  |
| Non-employed | 20(7.5) | 247(92.5) | 1 | 1 |
| Employed | 22(13.3) | 143(86.7) | 1.90(1.01,3.60)<sup>a</sup> | 2.58(1.25,5.37)<sup>b</sup> |
| **Perceived required time to reach health facilities** |  |  |  |  |
| ≥30 minutes | 11(6.1) | 169(93.9) | 1 | 1 |
| <30 minutes | 31(12.3) | 221(87.7) | 2.155(1.05,4.41)<sup>a</sup> | 1.89(0.82,4.37) |
| **Pre-pregnancy utilization of contraception (n=432)** |  |  |  |  |
| Yes | 37(12.6) | 257(87.4) | 3.83(1.47,9.97)<sup>a</sup> | 3.96(1.43,10.99)<sup>b</sup> |
| No | 5(3.6) | 133(96.4) | 1 | 1 |
| **Knowledge on key pregnancy danger signs (n=432)** |  |  |  |  |
| Not-knowledgeable | 18(6.9) | 243(93.1) | 1 | 1 |
| Knowledgeable | 24(14) | 147(86) | 2.20(1.16,4.19)<sup>a</sup> | 1.66(0.77,3.68) |
| **Birth order** |  |  |  |  |
| First | 10(13.7) | 83(86.3) | 1 | 1 |
| Second | 10(10.5) | 85(89.5) | 0.74(0.29,1.88) | 0.73(0.23,2.33) |
| Third | 10(8.3) | 111(91.7) | 0.57(0.22,1.44) | 0.48(0.16,1.51) |
| **Four and above** | 12(8.4) | 131(91.6) | 0.57(0.24,1.41) | 0.63(0.20,1.93) |
| **Desire on pregnancy (n=432)** |  |  |  |  |
| **Not planned** | 4(4) | 95(96) | 1 | 1 |
| **Planned** | 38(11.4) | 295(88.6) | 3.06(1.06,8.79)<sup>a</sup> | 3.49(1.07,11.43)<sup>b</sup> |
| **Time for first ANC booking (n=432)** |  |  |  |  |
| **At or after 16 wks.** | 15(4.5) | 322(95.5) | 1 |  |
| **Before 16 wks.** | 27(28.4) | 68(71.6) | 4.77(2.47,9.21)<sup>a</sup> | 10.7(5.09,22.68)<sup>b</sup> |
| **BPCR (n=432)** |  |  |  |  |
| **Not-well prepared** | 20(6.6) | 281(93.4) | 1 | 1 |
| **Well Prepared** | 22(16.8) | 109(83.2) | 2.84(1.48,5.40)<sup>a</sup> | 2.93(1.41,6.09)<sup>b</sup> |

COR=Crude Odds Ratio ,AOR=Adjusted odds ratio <sup>a</sup> significant in bivariate analysis at p-value of less than 0.25, <sup>b</sup> statistically significant association at p-value of less than 0.05 after adjusting for all the other variables in the model- BP/CR, time for ANC initiation, desire on pregnancy, perceived required time to reach health facilities, women's employment status, women's education status, pre-pregnancy family planning utilization, knowledge on key pregnancy danger signs, and birth order.

## DISCUSSION

In this study the level of continuum of care was 9.7%[95% CI: 6.9%, 12.5%] which is consistent with findings from study in Tanzania (10%)[17] but higher than study conducted in Cambodia (5%)[23], and lower than studies conducted in Pakistan (27%) and Egypt (50.3%) [21, 22]. The possible reason in this study might be due to time variation, variation in study design, and variation in source of data.

Women’s employment status was associated with completion of continuum of care for maternal and newborn health services. This finding is in line with prior study conducted in Egypt[21]. In contrast, this finding contradicts with that of previous study conducted in Pakistan [22]. In Pakistan, secondary data analysis found that being employed would decrease the chance of completion of continuum of care by 19%. This study found that the odds of completion along continuum of care were 2.586 times higher for employed women when compared to unemployed women. The discrepancy between two studies might be due to the difference in proportion of employed women. The possible reason might be due to the fact that employed women have more autonomy to decide on health care seeking because in this study the percentage of autonomy was 79% while 43.7% in Pakistan [21]. The percentage of autonomy for employed women was 86.1% when compared to 75.7% among non-employed women. This finding is supported by study from Bale Zone, South East Ethiopia which found that employed women were 5 times more likely autonomous when compared to their counterparts[29]. Autonomy in household decisions to seek health care, even if it did not show association with completion of CoC in this study, a number of studies found that it increases the probability of completion along continuum of care for MNH services. For instance, a 20% increase in Pakistan and 45% increase in south Asia and Sub-Saharan African countries was documented in previous studies [22, 30]. The other implication to this finding is that working women might have more access to gain money and plan for health care unlike housewives who solely engage in household duties.

Time for antenatal care initiation was one of the factors that facilitate completion of continuum of care in this study. This is plausible from the fact that women who present early for antenatal care are having of higher chance to complete the recommended optimum ANC visits and thus complete the first continuum required for completion of CoC. WHO also recommends on early antenatal care booking in order to women’s to attain adequate antenatal care visits and improve their health[31]. Besides that, in this study early antenatal booking is not only increase adequate antenatal care visit but also it helps to women’s to complete CoC. The reason for this finding is might be early booking is opportunity for health promotion and a powerful predictor of the content of antenatal care services [25,32,33]. In this study among women’s who booked early for antenatal care, around two third (64.2%) received full essential antenatal care services and more than two thirds (68.4%) attended antenatal care four or more times. Multiple studies supported the association between early ANC booking and quality ANC service. For instance, a study in Pakistan found that an early booking for antenatal care increases the probability of receiving the WHO recommended essential services during the pregnancy period when compared to late booking for antenatal care [32]. In India, women who booked early were 22 times more among women who had utilized minimum recommended antenatal care services(tetanus toxoid injection 2+, iron provision(100+)) when compared with women who booked late[25]. Thus, the association between early ANC booking and completion of CoC for MNH services in this study might be attributable to its effect on women’s access to quality service and adequate visits.. Furthermore, in this study among women’s who booked early, (62.1%) were informed about key danger signs during pregnancy. Women who attended ANC early were more likely to be better informed about pregnancy danger signs and obstetric complications[34, 35] and this may have effect on the subsequent health service utilization. This is supported by prior study conducted in Tanzania which documented a higher percentage (91%) of subsequent service utilization among women who recognized pregnancy danger signs earlier in their pregnancy[36]. A possible explanation would also include that pregnant women-attending ANC clinics early might have the chance to adapt to the health facility environment and this in turn would have helped them to avoid unnecessary fear and stress related to institutional service use. Moreover, early booking might help women to set birth plans in consultation with the ANC provider and hence increase women’s frequency of antenatal care visit, delivery and postnatal service use.

Another finding is the association between contraceptive utilization before recent birth and completion of continuum of care for maternal and newborn health service. This finding was supported by study conducted in Bangladesh where the odds of having SBA were 6.8 times higher for women’s reporting antenatal care visit four times and did used pre-pregnancy contraception when compared to women’s reporting both no ANC and pre-pregnancy contraceptive use [11]. The reason behind this might be women’s who utilized pre-pregnancy contraception were well informed about subsequent maternal and newborn services and set plans with health professional for next services. This was supported by findings from a systematic review and meta-analysis on preconception care where the odds of attending antenatal care for women who were counseled about subsequent services during preconception period were 39% higher when compared to their counterparts[37]. Similarly, it might also increase early entry into prenatal care[38]. In this study, two thirds of women’s who utilized pre-pregnancy contraception planned for pregnancy when compared to those who did not utilized contraception before their recent birth. This might have affected its association with completion along continuum of care for maternal and newborn health services utilization.

Birth preparedness and complication readiness is one of factors positively associated with completion of continuum of care. The odds of completion along continuum of care were higher among women’s who were prepared for birth and ready for complication as compared with their counterparts. This finding is a novel addition of this study and has a policy implication as BPCR is one of twelve WHO recommendations for health promotion in order to increase the use of skilled care at birth and to increase the timely use of facility care for obstetric and newborn complication during postnatal period[39]. Women who were well prepared for birth and its complications were likely to be educated, from socio-economic households and having a better knowledge on pregnancy, labor and delivery and postnatal newborn and maternal danger signs and thus in favor to complete the CoC for MNH services [27, 28, 40, 41]. Despite its greater implication in this study, 62(56.9%) of women who had ANC4+, 108(53.5%) who gave birth at health facilities and 107(60.8%) of who attended postnatal checkup were with less prepared for birth and its complications. A previous study in the same setting has also showed a lower practice of BPCR among pregnant women[41]. This results call for a further attention on improvements of women knowledge and practice on BPCR.

The odds of completion of continuum of care among women who planned pregnancy was higher when compared to their counterparts in the present study. The finding is inconsistent with findings from a study conducted in Ghana in 2015 [18]. This might be due to the difference in proportion of unplanned pregnancy between the two studies (22% in this study and 32.1% in the study from Ghana). A related study conducted in Ghana in 2018 found no association between pregnancy intention and completion along continuum of care for maternal and newborn health services[19]. The possible reason might be having of planned pregnancy reduce delay in prenatal care which might have increase the chance of frequent visits along continuum of care because in this study early antenatal care booking was 17.8% among women’s with planned pregnancy while 4.2% among women’s with unplanned pregnancy. This is supported by prior studies which showed that unplanned pregnancy associated with late initiation of antenatal care visit [42, 43]. In this study, almost all unmarried women’s pregnancies were unplanned, so that they may initially attempt to deny their pregnancies to themselves and to conceal them from others because most of the time premarital pregnancies were highly stigmatized [44]. As the result, women may become less motivated to seek care along continuum of care compared to women with planned pregnancy. This is supported by study conducted in one of districts in northern Ghana which found that the odds of completion along continuum of care among single women were 87% less likely compared to their married counterparts[18]. The study was not free of limitations. To mention, first, the study participants were women who gave birth the last 1 year. Given that they were asked to recall their pregnancy experience as far back as to 1.5 years, recall bias was possible. Second, the study was based on self-reports and this might have introduced social desirability bias. Moreover, the sampling frame was based on a secondary data which may not have had registered all the eligible women for the study. Thus, the findings of this study should be interpreted with consideration of these limitations.

## CONCLUSION

In this study, the coverage of completion along continuum of care was lower when compared to other studies. The main identified factors associated with completion along continuum of care were pre-pregnancy contraceptive utilization, pregnancy desire, birth preparedness and complication readiness, women’s employment status and timing for antenatal care booking. Despite their higher contribution to completion of continuum of care for maternal and newborn health services, the proportion of early booking of antenatal care and provision of essential selected elements of antenatal care services were very low.

## ACKNOWLEDGMENT

We would like to address sincere appreciation and warmest thanks to Arba Minch University, College of Medicine and Health Sciences, and Department of Public Health for provision of ethical clearance for the conduct of this study. We are also indebted to our data collectors, supervisors and the study participants.

## Supporting information

S1Fig. Schematic presentation of the sampling procedure

S1data.The raw data supporting the findings of this article

## Authors’ contributions

DH: Conceived and designed the study, conducted data collection and analysis and wrote the draft manuscript. DH and EA: Conceived and designed the study, supervised data collection, performed the statistical analysis and wrote the first draft manuscript. MK: Assisted in the design of the study, supervised data collection, participated in data analysis and interpretation and preparation of the draft manuscript. AT, TF and NB: participated in data collection, analysis, interpretation and preparation of the draft manuscript. All authors read and approved the final manuscript.

